# Breast Density Analysis Using Digital Breast Tomosynthesis

**DOI:** 10.1101/2023.02.10.527911

**Authors:** John Heine, Erin E.E. Fowler, R. Jared Weinfurtner, Emma Hume, Shelley S. Tworoger

## Abstract

We evaluated an automated percentage of breast density (BD) technique (PD_a_) with digital breast tomosynthesis (DBT) data. The approach is based on the wavelet expansion followed by analyzing signal dependent noise. Several measures were investigated as risk factors: normalized volumetric; total dense volume; average of the DBT slices (slice-mean); a two-dimensional (2D) metric applied to the synthetic images; and the mean and standard deviations of the pixel values. Volumetric measures were derived theoretically, and PD_a_ was modeled as a function of compressed breast thickness. An alternative method for constructing synthetic 2D mammograms was investigated using the volume results. A matched case-control study (n = 426 pairs) was analyzed. Conditional logistic regression modeling, controlling body mass index and ethnicity, was used to estimate odds ratios (ORs) for each measure with 95% confidence intervals provided parenthetically.

There were several significant findings: volumetric measure [OR = 1.43 (1.18, 1.72)], which produced an identical OR as the slice-mean measure as predicted; [OR =1.44 (1.18, 1.75)] when applied to the synthetic images; and mean of the pixel values (volume or 2D synthetic) [ORs ∼ 1.31 (1.09, 1.57)]. PD_a_ was modeled as 2^nd^ degree polynomial (concave-down): its maximum value occurred at 0.41×(compressed breast thickness), which was similar across case-control groups, and was significant from this position [OR = 1.47 (1.21, 1.78)]. A standardized 2D synthetic image was produced, where each pixel value represents the percentage of BD above its location.

The significant findings indicate the validity of the technique. Derivations supported by empirical analyses produced a new synthetic 2D standardized image technique. Ancillary to the objectives, the results provide evidence for understanding the percentage of BD measure applied to 2D mammograms. Notwithstanding the findings, the study design provides a template for investigating other measures such as texture.

## 1. Introduction

Breast density is a significant and well accepted breast cancer risk factor assessed from mammograms [1–4]. Areas of increased breast density (i.e., the degree of bright tissue) correspond to tissue with greater x-ray attenuation properties, as observed in mammograms used for clinical purposes. Increasing areas of dense tissue correspond to increased risk. We have been investigating techniques that quantify both breast density [5–13] and more generalized measures that capture texture in the image [14–16] and Fourier domains [15, 17] in two-dimensional (2D) *conventional* mammograms. Breast density is one factor among others that could be considered in the development of breast cancer risk prediction models for clinical purposes. Such models could be used for developing personalized healthcare strategies, such as setting risk-modulated screening intervals or modalities, providing the accuracy permits [18, 19].

In the current clinical environment, there are several breast cancer risk models used for specific purposes [4, 20] for example: the Gail model is used to advise on chemoprevention for reducing risk; the Tyer-Cuzick and BRCAPRO models are used to predict the probability of a BRCA mutation but have been supplanted by the National Comprehensive Cancer Network guidelines; and the Tyrer-Cuzick, BRCAPRO, and Claus models are useful for determining if supplemental imaging with magnetic resonance might be beneficial. It is worth noting, the BRCAPRO, Tyrer-Cuzick and Gail models incorporate breast density. Recently, the American College of Radiology provided recommendations for supplemental imaging based on risk in conjunction with breast density [21]. At present for screening in many high-income countries, risk is based primarily on age [22]. Risk prediction methods that use some type of image derived information (simple modeling through deep learning) show that texture may be an important factor, yet it is not commonly used in practice [4, 23, 24].

There are many methods under investigation for measuring both breast density and more generally texture [24–27]. The percentage of breast density measure (PD) has been studied for many years and has repeatedly shown to be significantly associated with breast cancer risk [28]. PD requires determining a threshold in a 2D mammogram; all pixels above this threshold are labeled as dense or otherwise labeled as non-dense creating a binary image. The final measure is derived from normalizing the dense area by the total breast area, presented as percentage. The Breast Imaging Reporting & Data System (BI-RADS) four state ordinal breast composition classification [29] has also been used as a risk measure and shows similar risk prediction capability across studies [1]. These measures capture the volumetric tissue characteristics projected onto a plane. Newly derived image markers are often compared to PD as it has been considered as the de facto benchmark standard. For example, recent work shows that PD produces a risk measure marginally superior to a commercially available volumetric breast density product when studying 2D full field digital mammography (FFDM) images [30].

Breast screening recently has largely transitioned from FFDM to digital breast tomosynthesis (DBT). DBT provides a *three-dimensional* (3D) rendering of the breast via stacked 2D images (slices) derived from 2D projection images acquired over a limited angular range. Clinical DBT images result from heavy processing. There is little published work with measures derived from clinical DBT images for risk factor purposes at this time [24, 31]. It is reasonable to assume that a more precise measure of breast density would result from analyzing volumetric images in comparison with conventional 2D mammograms. Accordingly, recent work in comparing an automated volumetric measure from DBT with measures applied to 2D mammograms shows improvements in risk prediction capability [32]. Moreover, recent modeling using DBT data, incorporating various images features, illustrates it is possible to guide image care [23].

We previously developed an automated PD approach (PD_a,_) that was evaluated in studies with digitized film and FFDM images [8, 10, 33]. Here we will apply this method to DBT data. A matched case-control study is investigated with women that have DBT volume datasets (2D slices) and synthetic 2D mammograms (C-Views). There are four main study objectives. First, we investigate an algorithm modification used recently when studying relatively low-resolution digitized film mammograms [33] and describe this algorithm in more detail. Secondly, we show how to derive different breast density metrics from DBT volume slices and compare these measures with their 2D synthetic counterpart as breast cancer risk factors. Thirdly, PD is modeled as a function of compressed breast thickness. As a fourth objective, we provide evidence describing the nature of PD using the volume results. Here we show why the normalization by the total breast area used in the conventional PD measure (i.e., 2D application) may be important for its performance and provide a method of producing a normalized 2D synthetic image. We will focus on image analyses in tandem with the epidemiologic considerations. Mammographic imaging technology has clearly demonstrated its capability of advancing relatively rapidly over a short timespan (film, to FFDM, to DBT). Therefore, it is important to continue fundamental image analytics. This may enable algorithm modification more easily when the next innovation occurs.

## 2. Methods

### 2.1 Study data and imaging

This study is comprised of 426 matched case-control (CC) pairs of women with DBT datasets. To derive a larger CC dataset, we took the union of women from two prior CC studies (one studying 2D FFDM images and another studying DBT data) that had DBT datasets. Both CC studies used the same participant selection protocol. The selection criteria and study populations were described previously [16, 17]. Briefly, cases were women with first time pathologically verified unilateral breast cancer from two sources: 1) women attending the breast clinics at Moffitt Cancer Center diagnosed with breast cancer, and 2) attendees of surrounding area clinics sent to this Moffitt for either breast cancer treatment or diagnostic workup and found to have breast cancer. Controls were attendees of this center that never had breast cancer and verified by a two-year follow-up (i.e., no history of breast cancer). Controls were individually matched to cases on: age (± 2 years), hormone replacement therapy (HRT) usage and current duration (never users or not current users in the prior 2 years, current user ± 2 years duration), screening history (any prior screening with time since last screening <30 months, any prior screening >30 months before baseline, no prior screening), and mammography unit (described below). Cranial caudal orientation mammograms were used for this study to reduce pectoral interference in the automated analyses. The unaffected breast image(s) were used for cases (all study images were acquired before treatment) and the matching lateral breast for controls. Population characteristics are shown in Table 1.

**Table 1:** Population Characteristics: participant numbers (n) are broken down by case-control status and totals. Where applicable, distribution means and standard deviations (SDs) are provided. Matching factors are indicated by asterisks.

| Measure or Characteristic | p-value | Case n | Case mean, SD, or relative frequency | Control n | Control mean, SD, or relative frequency | Total n | Total mean, SD, or relative frequency |
| --- | --- | --- | --- | --- | --- | --- | --- |
| **Age\*** | 0.7787 | 426 | 58.2 (11.6) | 426 | 58.2 (11.6) | 852 | 58.2 (11.6) |
| **BMI** | <0.0001 | 426 | 29.6 (6.9) | 424 | 27.7 (6.3) | 850 | 28.7 (6.7) |
| **Screening Group\*** | N/A |  |  |  |  |  |  |
| Group 1 |  | 262 | 61.50% | 262 | 61.50% | 524 | 61.50% |
| Group 2 |  | 87 | 20.42% | 87 | 20.42% | 174 | 20.42% |
| Group 3 |  | 77 | 18.08% | 77 | 18.08% | 154 | 18.08% |
| **HRT Usage\*** | N/A |  |  |  |  |  |  |
| Current |  | 36 | 8.45% | 36 | 8.45% | 72 | 8.45% |
| Not Currently |  | 390 | 91.55% | 390 | 91.55% | 780 | 91.55% |
| **Race** |  |  |  |  |  |  |  |
| Caucasian | 0.5394 | 340 | 79.81% | 348 | 81.69% | 688 | 80.75% |
| African American | 1.0000 | 47 | 11.03% | 46 | 10.80% | 93 | 10.92% |
| Asian | 0.5716 | 16 | 3.76% | 12 | 2.82% | 28 | 3.28% |
| Other | 1.0000 | 12 | 2.82% | 12 | 2.82% | 24 | 2.82% |
| More than one | 0.0703 | 7 | 1.64% | 1 | 0.23% | 8 | 0.94% |
| Unknown | 0.5488 | 4 | 0.94% | 7 | 1.64% | 11 | 1.29% |
| **Ethnicity** | <0.0001 |  |  |  |  |  |  |
| Non-Hispanic |  | 369 | 86.62% | 323 | 75.82% | 692 | 81.22% |
| Hispanic |  | 57 | 13.38% | 98 | 23.00% | 155 | 18.19% |
| Unknown |  | 0 | 0.00% | 5 | 1.17% | 5 | 0.59% |
| **Menopausal Status** |  |  |  |  |  |  |  |
| Premenopausal | 1.0000 | 114 | 26.70% | 111 | 26.06% | 225 | 26.41% |
| Postmenopausal | 0.7877 | 311 | 73.13% | 311 | 73.00% | 622 | 73.00% |
| Unknown | 0.3750 | 1 | 0.17% | 4 | 0.94% | 5 | 0.59% |

Image datasets were acquired with Hologic Dimensions DBT units (Hologic, Inc., Bedford, MA): *volume* images (in-plane 100μm average pixel spacing, 1mm slice thickness, with 10-bit pixel dynamic range); and synthetic (i.e., C-View) 2D images (100μm average pixel spacing with 10-bit pixel dynamic range). The in-plane pixel spacing varies for DBT *volume* slices and synthetic 2D images simultaneously and in unison for each acquisition, ranging from approximately 85μm to 106μm (about 100μm average) for our data. The number of slices in each DBT volume is approximately one slice per mm of compressed breast thickness.

### 2.2 Automated PD for conventional two-dimensional applications

Our automated PD detection mechanism (i.e., PD_a_) operates by analyzing signal dependent noise locally in standard 2D images (i.e., FFDM and digitized film images). We previously identified conditions that should be present for the algorithm to function *optimally* [10], that were refined here. We use the following idealization to explain the fundamental assumptions supporting the technique. We assume mammograms are captured with a linear detection technology that can be modeled as a cascaded Poisson process (pixel intensity scaled and/or bias shifted). This indicates that the variance and mean are linearly related. If we acquired the same image repeatedly and investigated the signal at a particular pixel (neglecting system blurring), this mean-variance relationship would approximately materialize. We have shown this approximates the case (as expected) for 2D raw (*for processing*) images acquired with either GE (100μm pitch with 14-bit dynamic range) [34] or Hologic Selenia units (70μm pitch with 14-bit dynamic range) [10] with an algorithm that requires a single mammogram realization to estimate the signal dependent noise functional form [34], rather than under controlled conditions with phantom imaging. For reference, this technique estimates the local variance as a function of the local mean for an arbitrary mammogram as briefly described and illustrated here. In practice, we do not acquire the same mammogram repeatedly in close time proximity. For the single acquisition case, we switch a time average with a spatial average to analyze locally varying signal dependent noise for the density detection. That is, if we consider that a mammogram varies slowly spatially, the statistical characteristics in small n × n pixel region can approximate the same characteristics of the signal at an isolated pixel after n^2^ acquisitions. In images where the pixel representation has larger values for dense tissue and smaller values for adipose tissue (i.e., monochrome 1 format or mammograms used for clinical purposes), both the local variance and mean will be greater in a small dense tissue regions (fibro-glandular) in comparison with these quantities from small non-dense (adipose) regions.

More specifically, the PD_a_ algorithm is two-stage detection technique that determines a global reference variance for adipose tissue (over the entire breast area) in a high pass filtered version of the image. In the first detection stage, this reference variance is used to make comparisons with local variances across the breast area. Local regions that deviate significantly from this reference are labeled as dense or otherwise non-dense by default. In the second detection stage, the reference variance is refined by estimating it from non-dense areas determined in the first stage. The localized comparisons are then repeated with the refined reference resulting in the PD_a_ labeled image. Each stage requires a threshold based on a significance level selected a priori and empirically from a Chi-square distribution. In applications, such as digitized film or applying the method to other images that have been processed in some form, the approximate linear form between mean and noise variance as estimated from FFDM data can take on a nonlinear relationship. Our explorations have shown (1) the near linear relationship in *for processing* images does not offer enough contrast to determine references for the adipose variation to label the images optimally, (2) applying a nonlinear transform to the image before deploying PD_a_ can have a beneficial impact on the algorithm [10, 14, 35], and (3) if the noise has been reduced in the image by some prior processing, it could reduce algorithm performance as explained below.

The mechanics of the PD_a_ method are outlined to facilitate the modification description. The algorithm operates by first applying a 2D wavelet filter (i.e, filter bank) to the image, *isolating* the noise. Here, we use noise to define *high spatial frequency chatter* constrained to the breast area. The filter output takes the form of an image expansion [36, 37] expressed as

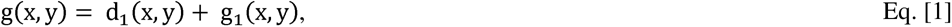

where g(x,y) is the image, d_1_(x,y) and g_1_(x, y) are the 2D high and low half bandpass filtered output images respectively, and (x,y) are pixel coordinates. Eq. [1], although complete, is a truncated one level expansion. Using the wavelet orthogonality conditions, the variance of g(x,y) can be expressed as

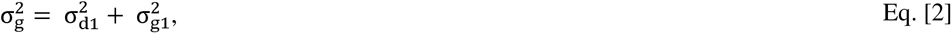

where the variances on the right side of Eq. [2] are the components corresponding to the terms on the right side of Eq. [1], respectively. Equation [2] holds globally and locally to a good approximation. The adipose reference values and density detection are established in d_1_(x,y), and g_1_(x,y) is not used (for PD_a_). In the past when applying the method to digitized film or different FFDM technologies, we have not found it necessary to adjust the filter’s bandpass characteristics when the pitch or pixel spacing varied. As reference, when characterizing signal dependent noise relationships, we use the same expansion in Eq. [1] but both components are analyzed; the local noise variances are determined in d_1_(x,y) and the respective local mean signals in g_1_(x,y) as the mean of d_1_(x,y) is zero. When moving g_1_(x,y) to the left side of Eq. [1], in images with smaller pitch, d_1_(x,y) often resembles *residu*e because the power spectral properties of (2D conventional) mammograms follow an inverse power law [5, 38, 39] (i.e., the image variance is well approximated by 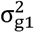 in Eq. [2]). If g(x,y) has been processed (prior to applying PDa) in such a way as to reduce the noise variance (i.e., some form of low pass-filter operation, or an operation that increased the pixel spacing), the algorithm may perform less than optimally because 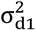 may become parasitic or *nonexistent*. The empirical correction [33] that can enhance the performance results from multiplying g(x,y) in Eq. [1] by a random noise field given by

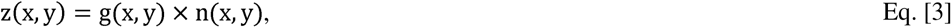

where n(x,y) is a normally distributed zero-mean, unit variance, noise. Equation. [3] is referred to as multiplicative noise and has similarities to models used in in ultrasound [40], Synthetic Radar Aperture images [41] and generally for characterizing signal dependent noise [42], where g(x,y) is the uncorrupted signal. Here, we make the approximation that g(x,y) is the uncorrupted image. Applying the same wavelet expansion to a modified (*corrupted*) mammogram, z(x,y), in Eq. [3] gives

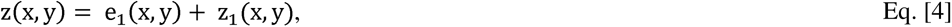

where e_1_(x,y) and z_1_(x,y) correspond to the components in Eq. [1], respectively, with the same interpretations relative to z(x,y). The expression for the variance for z(x,y) follows that of Eq. [2] giving

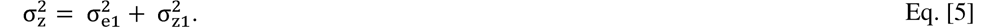

In contrast in Eq. [5], 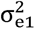 represents ¾ of the variance of z(x,y), which is a characteristic of white noise and as an approximation applies locally as well. To keep the presentation to an acceptable limit, we will show the variance components in Eq. [2] and [5] empirically here with examples. Briefly, the wavelet high half bandpass covers ¾ of the 2D frequency plane and the lower half bandpass covers ¼ of this plane. When expanding white noise in accord with Eq. [1], these variance proportions apply in each band respectively. We use Eq. [4], as the modification for this report, which was applied to all mammograms before applying PD_a_ except when noted. A different noise field realization is used for each mammogram when applying Eq. [4]. We used a search box with n×n = 4×4 pixels as the standard size. This creates an image representing the local variances (local variance map). Wavelet expansion images as well as the variance images are used for illustrations. Through experimentation using Eq. [4] (not shown) with 2D *for presentation* images from General Electric Senographe 2000D (Milwaukee, WI) [180 case-control pairs] and Hologic Selenia units (320 case-control pairs), we found that a 0.1 significance level was robust for both detection stages (i.e., determined from images that were processed in some way after the detection process). As part of this investigation (verification), detection thresholds (both stages) were set using a significance level = 0.1 with 15 degrees of freedom (i.e., n^2^-1) without analyzing DBT data with the modified methodology a priori.

The merits of equations 1-5 will be analyzed with examples of g(x,y) and z(x,y). Two study participants will be selected at random. Their images will be for illustrations and example analyses. For DBT illustrations, we will use the central slice images. We will also show the signal dependent noise plots derived from the g(x,y) and z(x,y) images from Eq. [3] and [4], respectively. The variances from Eq. [2] and Eq. [5] will also be evaluated. As a control example, we expanded a white noise field using the Eq. [1] and investigated the component variances as comparators. To make some illustrations clear for viewing purposes, we have excised the largest rectangle that can fit within the breast image that has a small margin on the breast perimeter side excluded, referred to as the largest rectangle below; the algorithm was described in detail elsewhere [17].

### 2.3 Breast Density Measurement Modeling from DBT

Multiple breast density measures can be derived from PD_a_ when applied to DBT datasets. We use a particular subscript convention to define the image format where PD_a_ was deployed, noting it is the same algorithm: (1) PD_syn_ when applied to synthetic 2D images, and (2) PD_s_ when applied to DBT slices. At the local region within a slice, we assume that the density detection process approximates the probability that the region has the x-ray attenuation coefficient of dense breast tissue, and it makes the decision based on this probability (i.e., referred to as the probability conjecture below).

For the volume derivations, we let a given DBT volume image have N slices with an isotropic thickness, t, measured in mm and pixel area given by A measured in mm^2^. The i^th^ slice has n_i_ pixels in the breast area with d_i_ pixels labeled as dense from PD_s_ (or generally, by any other method of applying PD). The expression for the breast volume (BV), required in the derivations, is given by

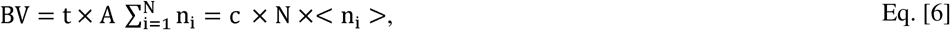

where the brackets indicate the average (expectation) operator, < n_i_ > is the average number of pixels over the slice breast areas, and c = txA As a measure, the total dense tissue volume (D_v_) within the breast volume is given by

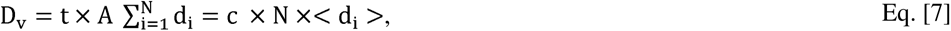

where <d_i_> is average of the N-PD_s_ findings. Using the BV and D_v_ expressions, the volumetric PD measure (PD_v_) is given by

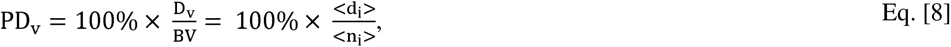

Another metric can be developed by applying PD_s_ to each slice and taking the mean (PD_m_) giving

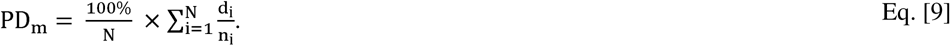

Equation [9] is an indication of why the normalization for PD may be important, as follows. Making the approximation that the breast area in each slice is constant with n pixels gives

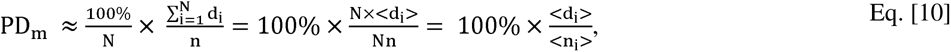

where n = <n_i_>. Then, when the breast areas are similar in each slice, PD_v_ ∼ PD_m_; this approximation will be evaluated. After applying PD_s_ to each slice, the labeled volume slices can be projected (summed) onto a plane giving a (coarse) 2D image with pixel values ranging between zero and N. The summation of pixel values within this image gives the projected total (PT) expressed as

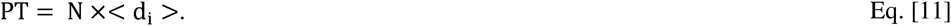

If we assume the traditional PD thresholding in 2D captures the dense pixel proportion above a given location within the compressed breast, the total number of dense pixels in a 2D labeled image is approximately the normalized projected total (NPT) determined by dividing Eq. [11] by N giving

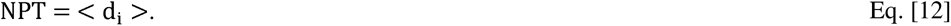

Given there are n pixels within the breast area, the standardized 2D measure is approximated by

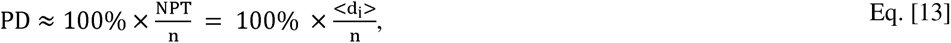

which is just Eq. [8] or Eq. [10] relabeled. The projected image normalized by N and multiplied by 100% produces a standardized synthetic image [s(x,y)], where each pixel represents the percentage of dense volume above its location; we note, these interpretations follow from the probability conjecture defined above. In the above developments, the image parameters (A or t) were not relevant except for D_v_. When making assessing D_v_ across women, t is constant while A will vary and must be accounted for in the metric.

We will investigate the breast cancer association provided by PD_syn_ (with and without noise multiplication), PD_vol_ and PD_m_. We will also investigate the associations from the isolated center DBT slice (i.e., exploring the possibility that PD from one slice may be representative of the volume). As further comparators, we will evaluate the mean and standard deviation of the pixel values within the DBT volume without applying PD_s_ processing, referred to as m_vol_ and v_vol_. Likewise, we will use the mean from the 2D synthetic images (m_syn_) as another comparator. All measures will be compared with those produced by PD_syn._(without noise multiplication).

To study breast density characteristics throughout the DBT volume from PD_s_, we used an empirically driven approach. We investigated a second-degree polynomial model

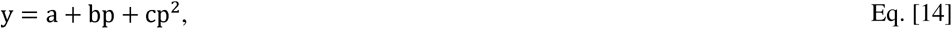

where y is PD_s_ for each slice, p is the normalized independent slice number variable ranging from [1, 100] measured in percentage [i.e., the closest PD_s_ measure to the breast support surface to the furthest measure from the support surface], and (a, b, c) are parameters to be determined with curve fitting analysis. The slice distance can be recovered approximately by 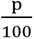 × compressed breast thickness. The convention used for increasing p follows that of the clinical volume rendering from the image header files. This normalization for distance puts the fit-parameters on the same scale over all participants. We investigated parameter distributions (empirical normalized histograms) and made comparisons across CC status. The normalized distance for the maximum PD_s_ quantity derived from Eq. [14] is given by

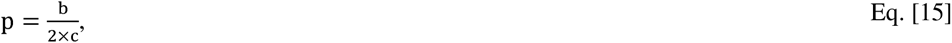

which is investigated below.

### 2.4 Statistical Analyses

Image measure associations with breast cancer were evaluated using conditional logistic regression while controlling for body mass index (BMI) and ethnicity. Unadjusted models are included in the tables for completeness. We used odds ratios (ORs) with 95% confidence intervals (CIs) as the primary metric to evaluate and compare breast cancer associations between the various image measures defined above. ORs were estimated for continuous measures in per standard deviation (SD) increment. We note, an OR derived from a CC study from a given population is often used as an approximation for relative risk for the same population, providing the disease incidence is small. Although building predictor models is not the purpose of this report, for completeness the area under the receiver operator characteristic curves (Azs) were calculated with 95% CIs to summarize the discriminatory ability for each model. Both ORs and Azs are presented with CIs parenthetically. Note, the matching design in these CC studies was implemented specifically to isolate the associations of image measures with breast cancer and generally precludes developing risk prediction models, which require detailed information regarding selection probabilities. Pearson correlation coefficient (R) was used to show the linear relations between select breast density measures and with BMI.

## 3. Results

### 3.1 Population characteristics

Table 1 shows the characteristics of the CC participants. As expected, matching variables (age, screening, HRT) were similar across CC groups. Similarly, neither race (Caucasian, African American, or Asian) or menopausal status varied significantly by status. The similarity of menopausal status is that it is likely a surrogate for age. In contrast, both ethnicity (Hispanic, non-Hispanic) and BMI (larger for cases) varied significancy. The BMI finding is expected, as increased risk is associated with increased BMI [43]. The difference in ethnicity is due to the shifting demographics at our clinics.

### 3.2 Illustrations and Related Analyses

Figure 1 (top row) shows the images from the two participants selected at random in the following order from left to right: (a) C-View image for illustration-1; (b) slice from the DBT volume for illustration-1; (c) C-View image for illustration-2; and (d) volume slice image from illustration-2. Illustration-1 has 89μm pixel spacing and illustration-2 has 107μm pixel spacing. Breast areas for illustration-1 are (a) C-View-78 cm^2^ and rectangle-34 cm^2^; (b) central slice-78 cm^2^ and rectangle-33.cm^2^; Breast areas for illustration-2 are: (c) C-View-173 cm^2^ and rectangle-77 cm^2^; (d) slice-170.cm^2^ and rectangle-75.cm^2^. The images in the first row and regions in the second row assume the position of g(x,y) in Eq. [2] and the regions in the third row assume the position of z(x,y) in Eq. [4]. The same ordering was used throughout the presentation in the figures. The largest rectangle is outlined in each image and the respective regions used for improved viewing purposes are shown in the second and third rows of Figure 1. Although the structure in the volume slices track that of the C-View images, it appears less resolved. Figure 2 shows the relevant wavelet expansion images (excised regions): in the top-row, d_1_(x,y) derived from g(x,y) in Eq.[1]; and in the bottom-row, e_1_(x,y) derived from z(x,y) Eq. [4] (bottom-row). These figures illustrate the chatter description of noise and show spatial variation; visual comparisons between like images in the top and bottom rows show little difference. Quantitatively, the differences are summarized in Table 2. These show that the d_1_(x,y) image accounts for less than 3% of the variances, whereas the e_1_(x,y) images account for approximately 75% of the variances as predicted (noting, the remaining portion of the variances is accounted for in the low half band pass filtered images). As a control, Figure 3 shows the expansion of white noise. In this example, about 74% of the variance was accounted for in d_1_(x,y). Thus, the z(x,y) images share a characteristic of white noise. In contrast, the random noise expansion images show uniform variation in texture, although differences in texture between d_1_(x,y) and g_1_(x,y) are clearly different as expected (chatter versus more coarse variation). Figure 4 shows the signal dependent noise plots. The top-row shows the plots derived from g(x,y) and the bottom from z(x,y); qualitatively, the plots do not appear to have common characteristics other than a non-linear form noticeable in the top row. Figure 5 shows the intermediate variance images (regions) resulting from z(x,y). There are two things to note from these illustrations: (1) the local nature of the detection (i.e., salt and pepper appearance); and (2) the density labeling tracks well with the areas that appear to be dense tissue in their counterpart images in Figure 1. Figure 6 shows the PD_syn_ and PD_s_ detection outputs for the illustrations: the top row shows the outputs when applied to g(x,y) and the bottom row when applied to z(x,y). A comparison between like images in the top and bottom rows indicates that the detection in the bottom row appears more resolved, although this is a subjective assessment. Figure 7 shows the standardized, s(x,y), images for these illustrations. These appear as processed images (*for presentation*) but with level contrast as they are 8-bit images with limited dynamic range. These illustrate another technique for creating synthesized 2D images in contrast to the C-View type images.

**Figure 1.**
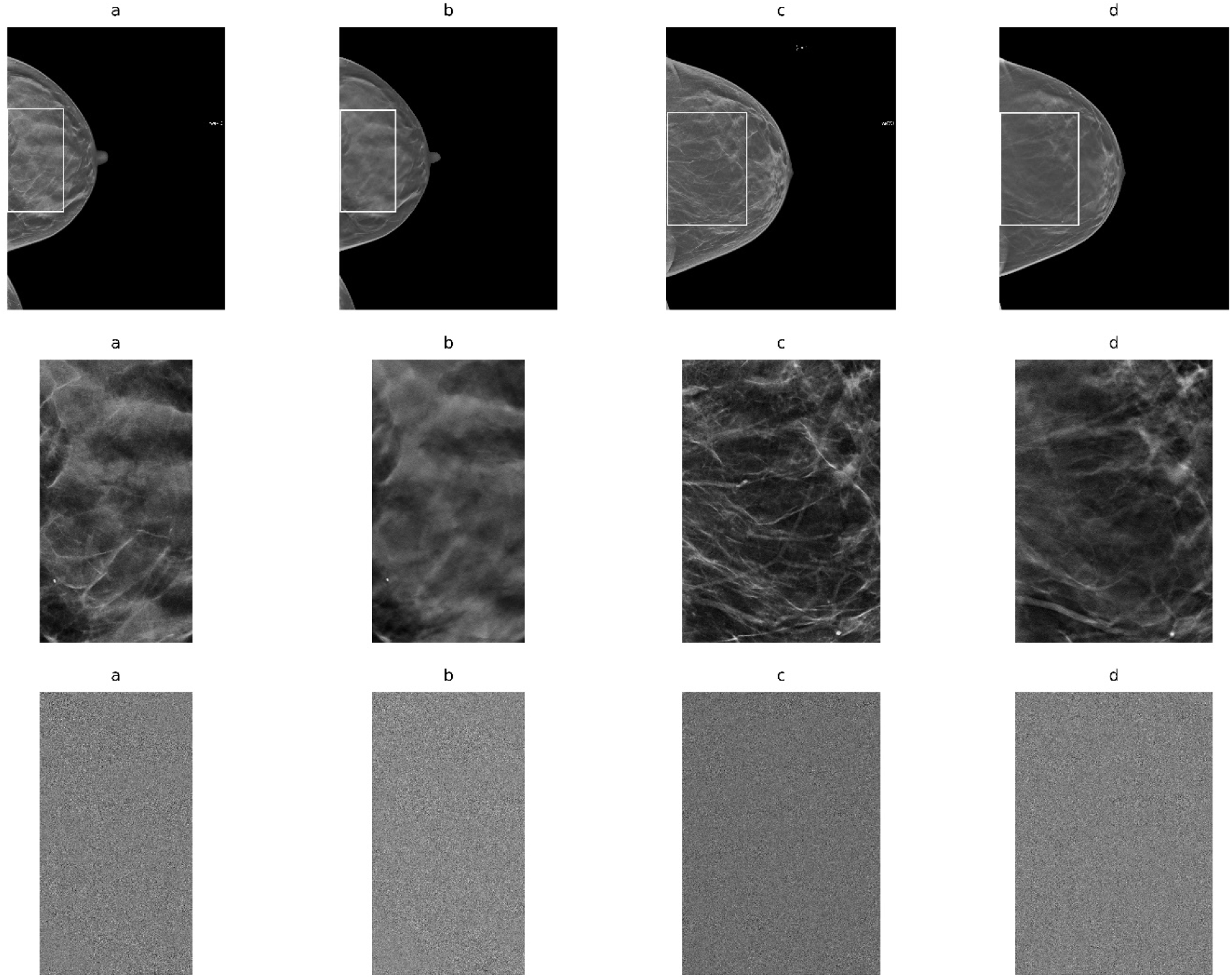
Image Illustrations: in the top row (a) C-View image for illustration-1; (b) respective central slice image; (c) C-View image for illustration-2; and (d) respective central slice image. Outlines in the top-row images are the largest rectangles that fit within the breast areas. These regions are shown in the second row with the same ordering for illustration purposes. The regions in the third row show the respective noise multiplication from Eq. [4]. Illustration-1 has 89μm pixel spacing and illustration-2 has 107μm pixel spacing. Breast areas for illustration-1 are (a) C-View-78 cm^2^ and rectangle-34 cm^2^; (b) central slice-78 cm^2^ and rectangle-33.cm^2^; Breast areas for illustration-2 are: (c) C-View-173 cm^2^ and rectangle-77 cm^2^; (d) slice-170.cm^2^ and rectangle-75.cm^2^.

**Figure 2.**
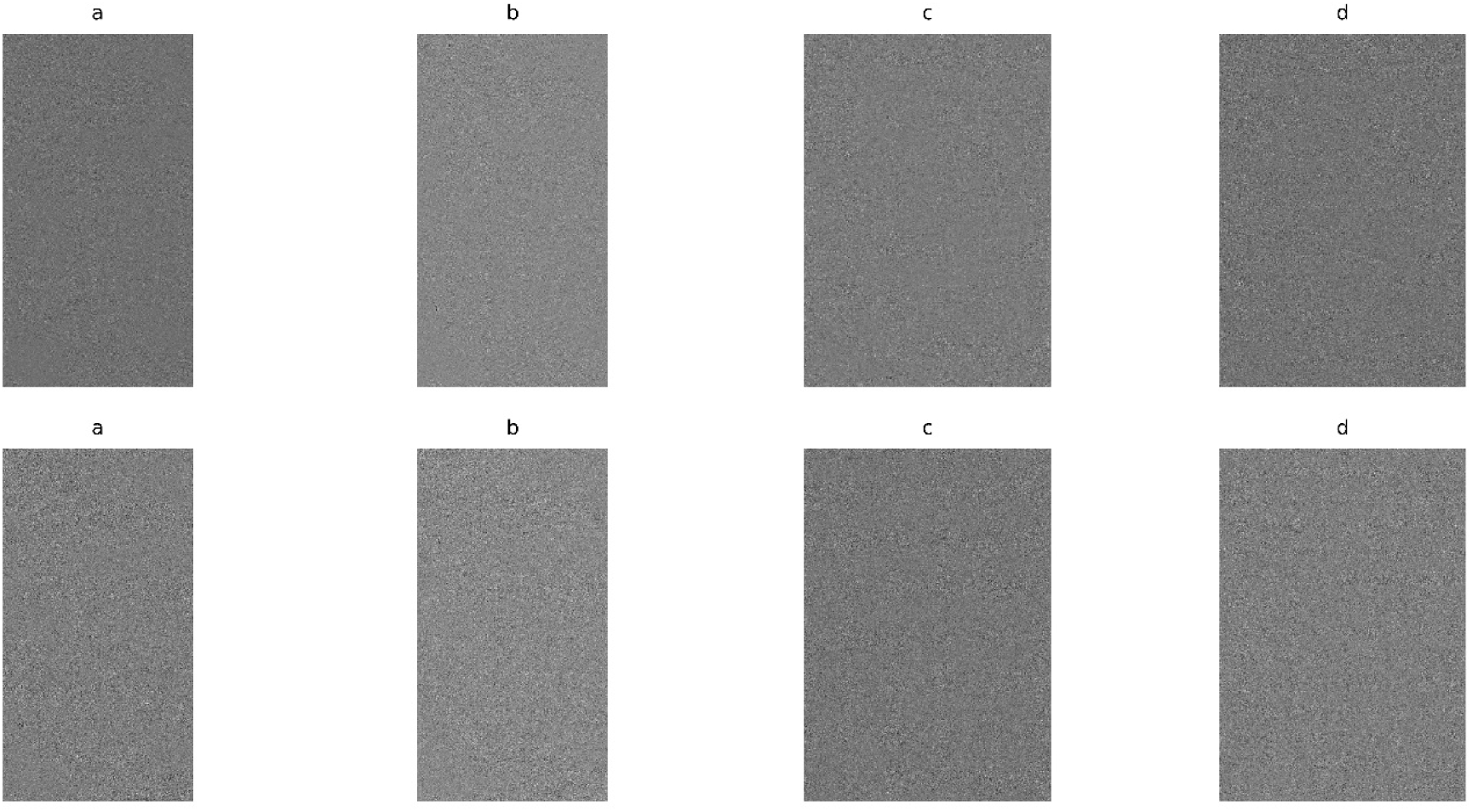
Wavelet Expansion Component Illustrations: these show the wavelet expansion components corresponding to the regions in the second row of Figure 1 excised from the illustration images. The top row shows the d_1_(x,y) components from Eq. [1]: (a) derived from the C-View image for illustration-1; (b) respective central slice image; (c) derived from the C-View image for illustration-2; and (d) respective central slice image. In the same order, the bottom row shows the respective e_1_(x,y) components (from the third row of Figure 1) derived from noise multiplication in Eq. [4].

**Figure 3.**
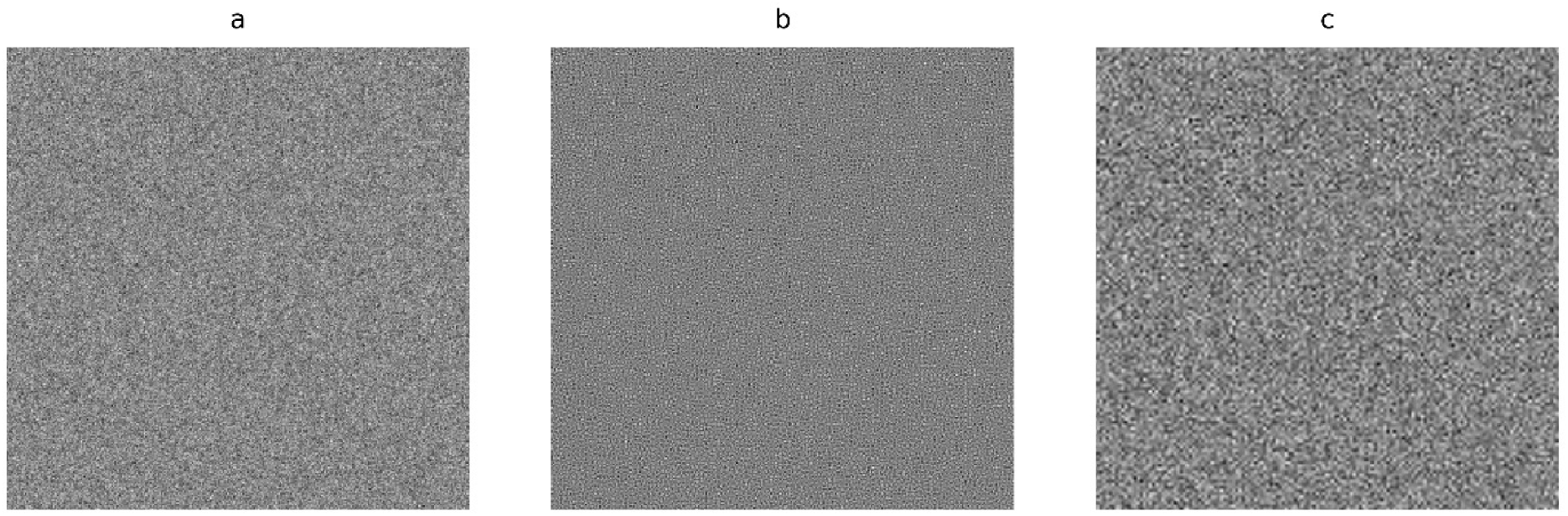
Wavelet Expansion of Random Noise: we use this as a control experiment to show the variances from the wavelet expansion images from Eq. [2] for noise and to describe what we have termed chatter: (a) g(x,y), random noise image; (b) d_1_(x,y), high pass expansion image (uniform chatter); (c) f_1_(x,y), low pass expansion image.

**Figure 4.**
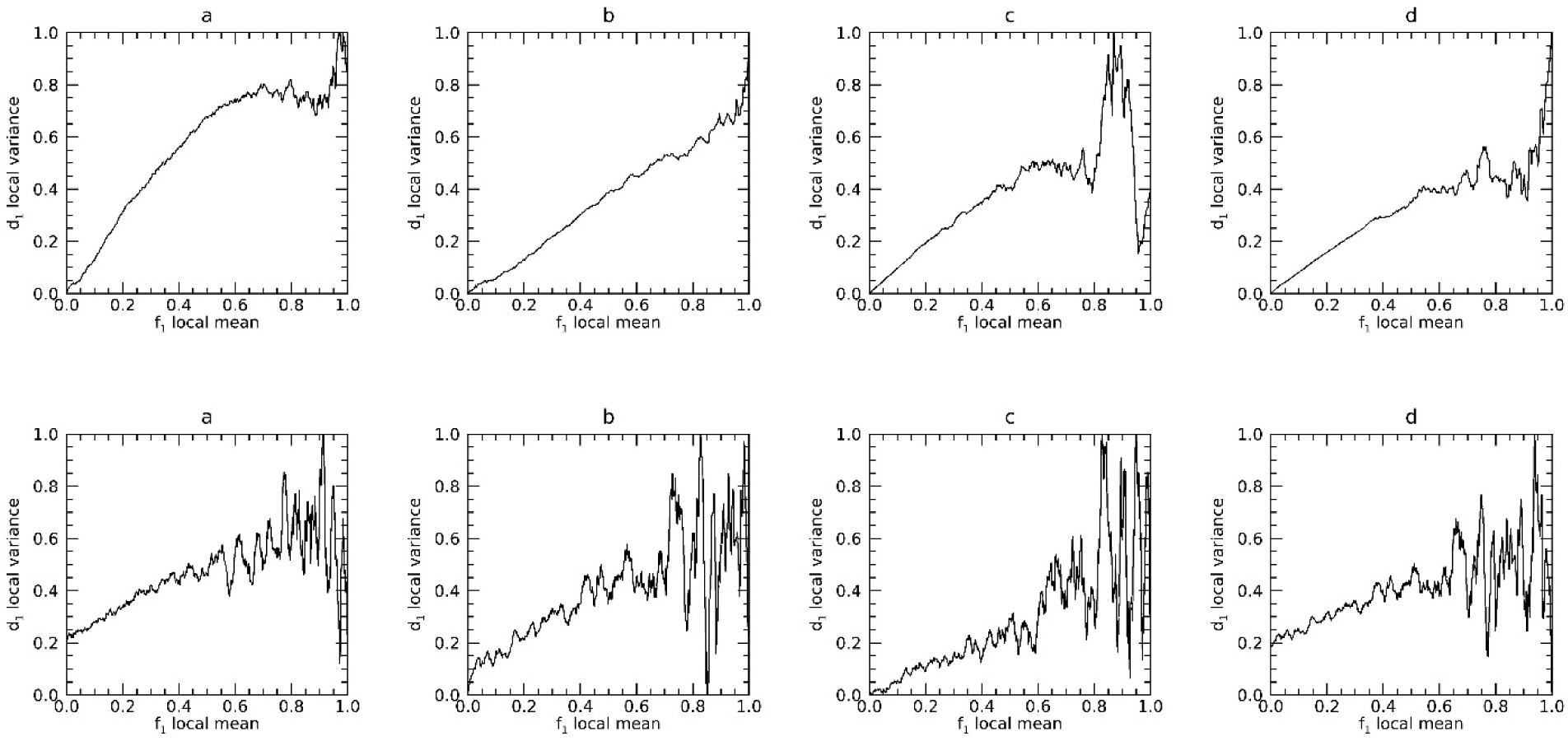
Signal Dependent Noise Plots: the top row shows the plots resulting from Eq. [1] (without noise multiplication) for the illustration images: top row, (a) derived from the C-view of illustration-1; (b) derived from the respective central slice image; (c) derived from the C-view illustration-2; and (d) derived from the respective central slice. The bottom row shows the respective plots derived from Eq. [4] (with noise multiplication) using the same ordering.

**Figure 5.**
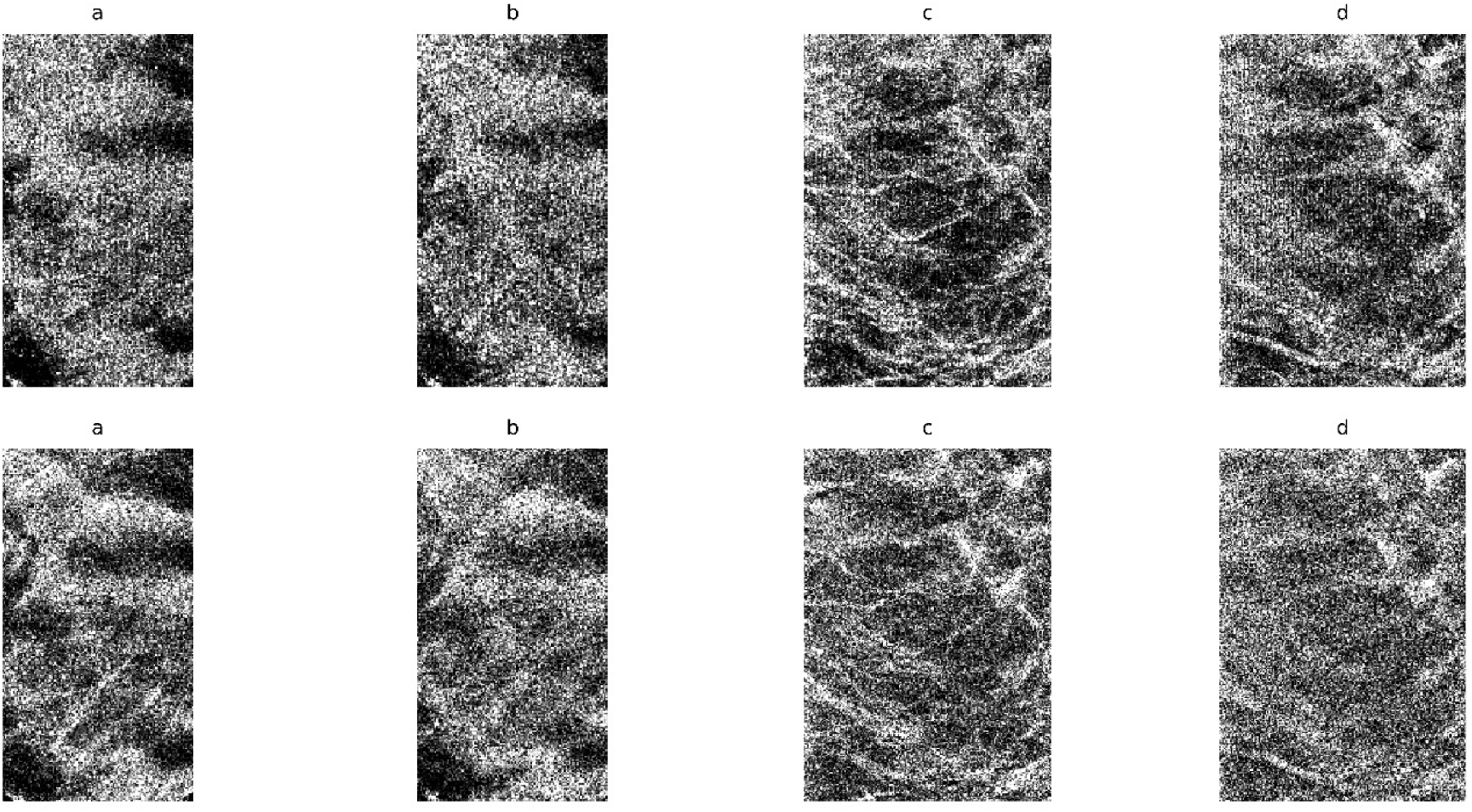
Variance Image Illustrations: these show the variance images for the regions shown in Figure 1, respectively. The top row results from Eq. [1] (without noise multiplication) and the bottom row from Eq. [4] (with noise multiplication): top row, (a) derived from the C-view of illustration-1; (b) derived from the respective central slice image; (c) derived from C-view of illustration 2; and (d) derived from the central slice. The bottom row is ordered as the top row. Window levels and widths are the same for all illustrations: set with the cumulative frequencies, where the lower 10% of the pixel values were set to the 10^th^ % value and the upper 10^th^ % percent of the pixel values were set to 90^th^ % value.

**Figure 6.**
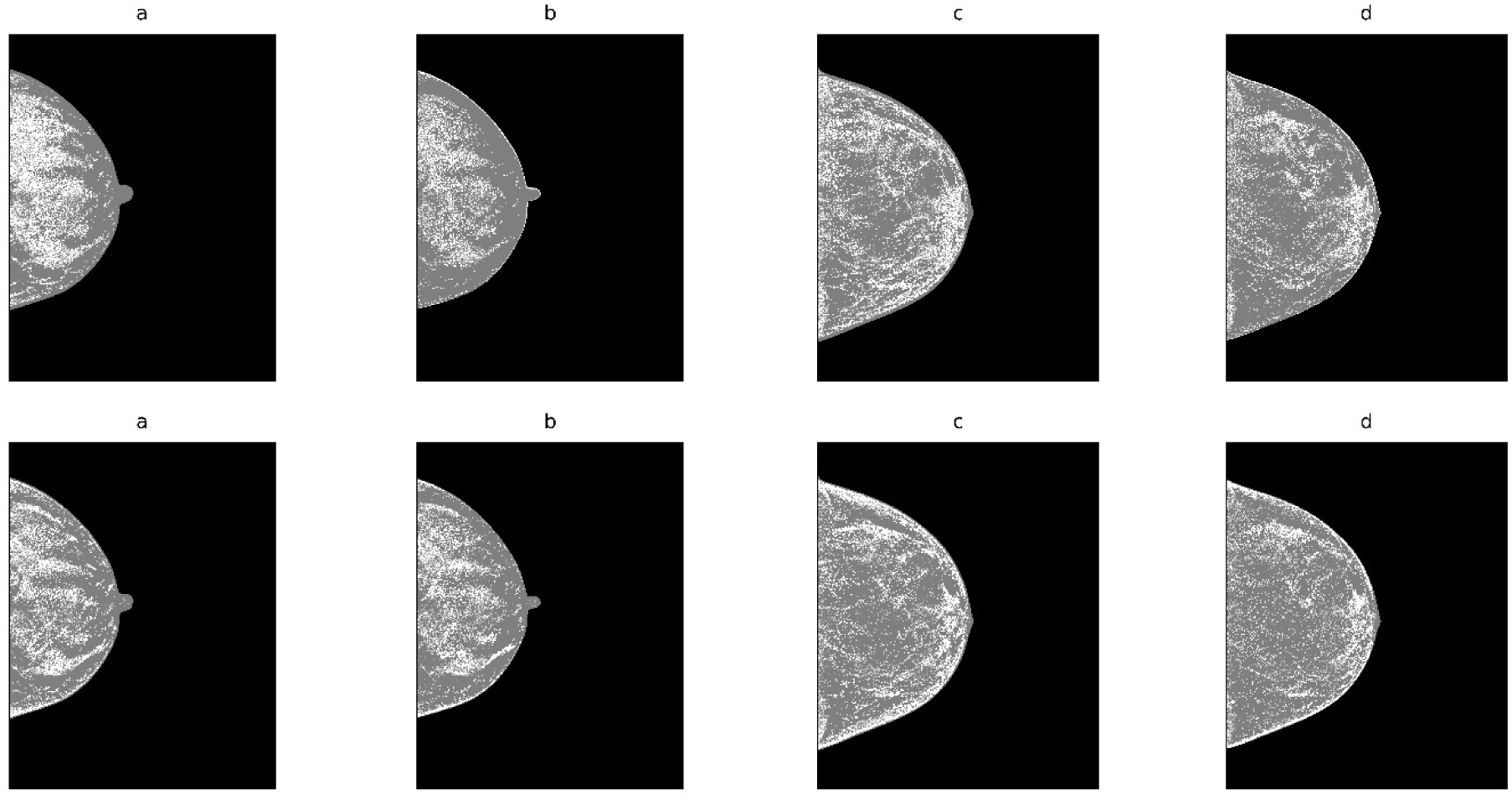
Breast Density Detection: the top row shows the detection resulting from Eq. [1] (without noise multiplication) in the top row, (a) C-view, illustration-1; (b) respective central slice image; (c) C-view, illustration-2; and (d) the respective central slice. The respective detection images resulting from Eq. [4] (with noise multiplication) are shown in the bottom row with the same ordering.

**Figure 7.**
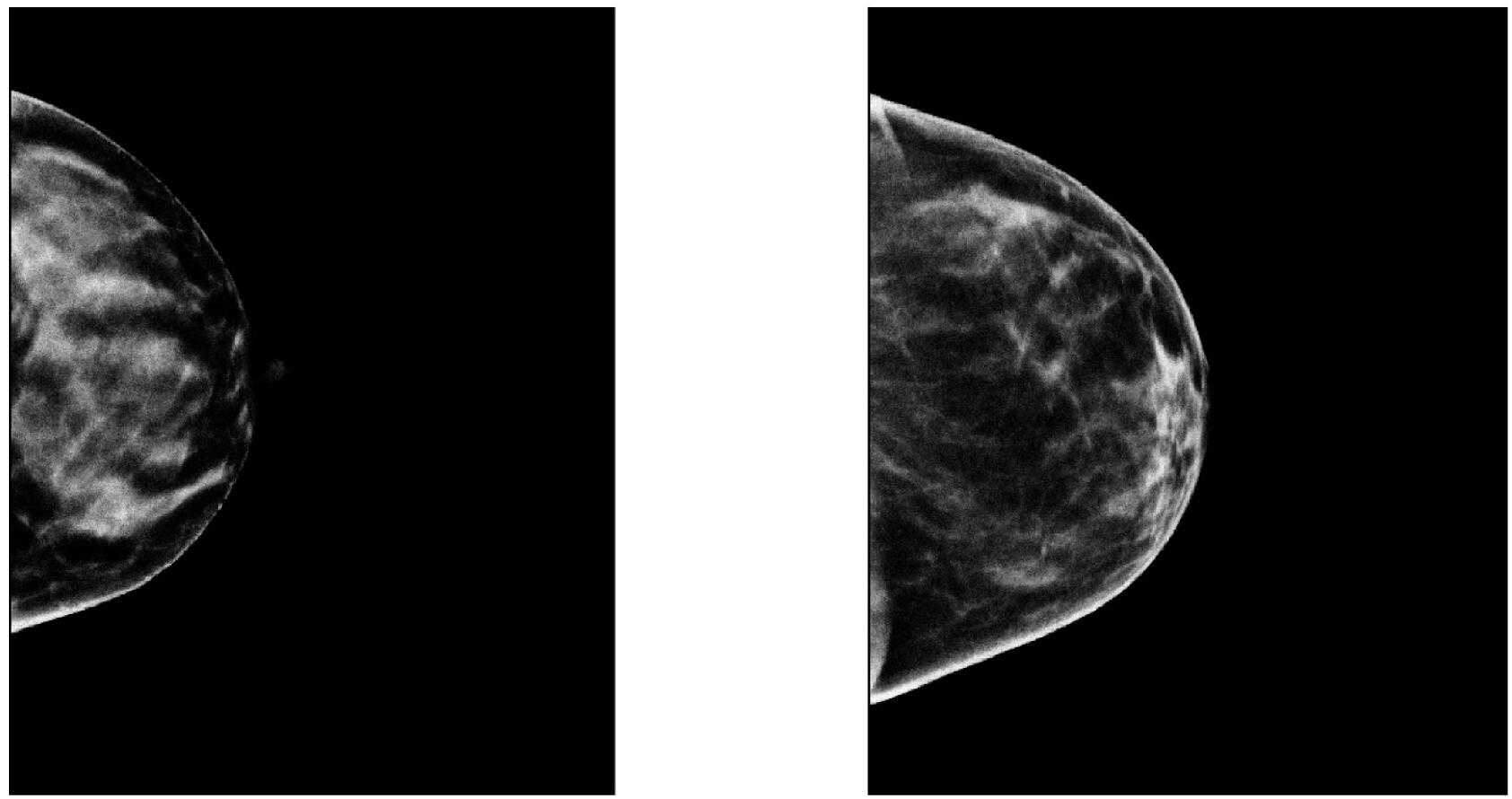
Projected Standardized Synthetic Breast Density Images: these show the standardized, s(x,y), images for illustration-1 (left) and illustration-2 (right) resulting from Eq. [4]. Pixel values represent the percentage of dense tissue above their locations.

**Table 2.** Variance Examples: these show the variances as a function of wavelet expansion images for both g(x,y) [g], and z(x,y) [z]: high pass-variances where calculated from d_1_(x.y) [d_1_] or e_1_(x,y) [e_1_], and the low-pass variances were calculated from g_1_(x,y) [g_1_] or z_1_(x,y) [z_1_], respectively. Percentages of the variance for these components are provided parenthetically.

|  | Illustration-1 |  |  | Illustration-2 |  |  |
| --- | --- | --- | --- | --- | --- | --- |
|  | g / z variance | d<sub>1</sub> / e<sub>1</sub> variance | g<sub>1</sub>/z<sub>1</sub> variance | g / z variance | d<sub>1</sub> / e<sub>1</sub> variance | g<sub>1</sub>/z<sub>1</sub> variance |
| C-view (g) | 5444.2 | 107.5 (1.97%) | 5329.1 (97.88%) | 2957.9 | 69.9 (2.34%) | 2882.4 (97.45%) |
| C-view (z) | 205041.5 | 153164.3 (74.70%) | 513070.8 (25.05%) | 158485.9 | 118203.1 (74.58%) | 39833.5 (25.13%) |
| Slice (g) | 3098.7 | 30.87 (1.00%) | 3063.7 (98.87%) | 1216.8 | 46.0 (3.78%) | 1165.8 (95.8%) |
| Slice (z) | 168340.3 | 125738.2 (74.69%) | 42134.6 (25.03%) | 136409.8 | 101801.9 (74.63%) | 34187.9 (25.06%) |

### 3.3 Measurement Modeling

Equations [8–10] show that PD_v_ and PD_m_ should be equivalent under the breast area similarity approximation. Figure 8 shows the scatter plots of these two measures (points) and the fitted regression line (solid red). Regression analysis gave R ≈ 1.0, slope ≈ 1.002, and intercept ≈ - 0.0201, indicating the two measures are essentially *identical*. This shows that both Eq. [13] and the interpretation of the synthesized images shown in Figure 7 are valid. DBT slice modeling using Eq. [14] is shown in Figure 9 for the illustrations. Note, the maximum density location occurs at p ≈ 0.41, and there is a cluster of PD_s_ quantities very similar to the maximum in close slice proximity in both illustrations. Figure 10 shows the histograms for the Eq. [14] coefficients separated by case-control status. Averaging like coefficients for cases and controls gave: (a, b, c)_mean_ ≈ (21.1, 0.06, -0.008) and (a, b, c)_mean_ ≈ (20.8, 0.07, -0.0008), respectively. Applying a t-test across groups showed only the intercept (i.e., a) varied marginally (p-value ≈ 0.046), as expected. Using parameter-means, the position with the greatest PD_s_ finding from Eq. [15] is approximately p ≈ 42 for either group. Empirically the mean maximum was p ≈ 40.7 with a standard error ≈ ± 0.28, showing close agreement with the model. Considering these findings, we investigated the breast cancer associations determined from the slice with the largest PD_s_ finding as well.

**Figure 8.**
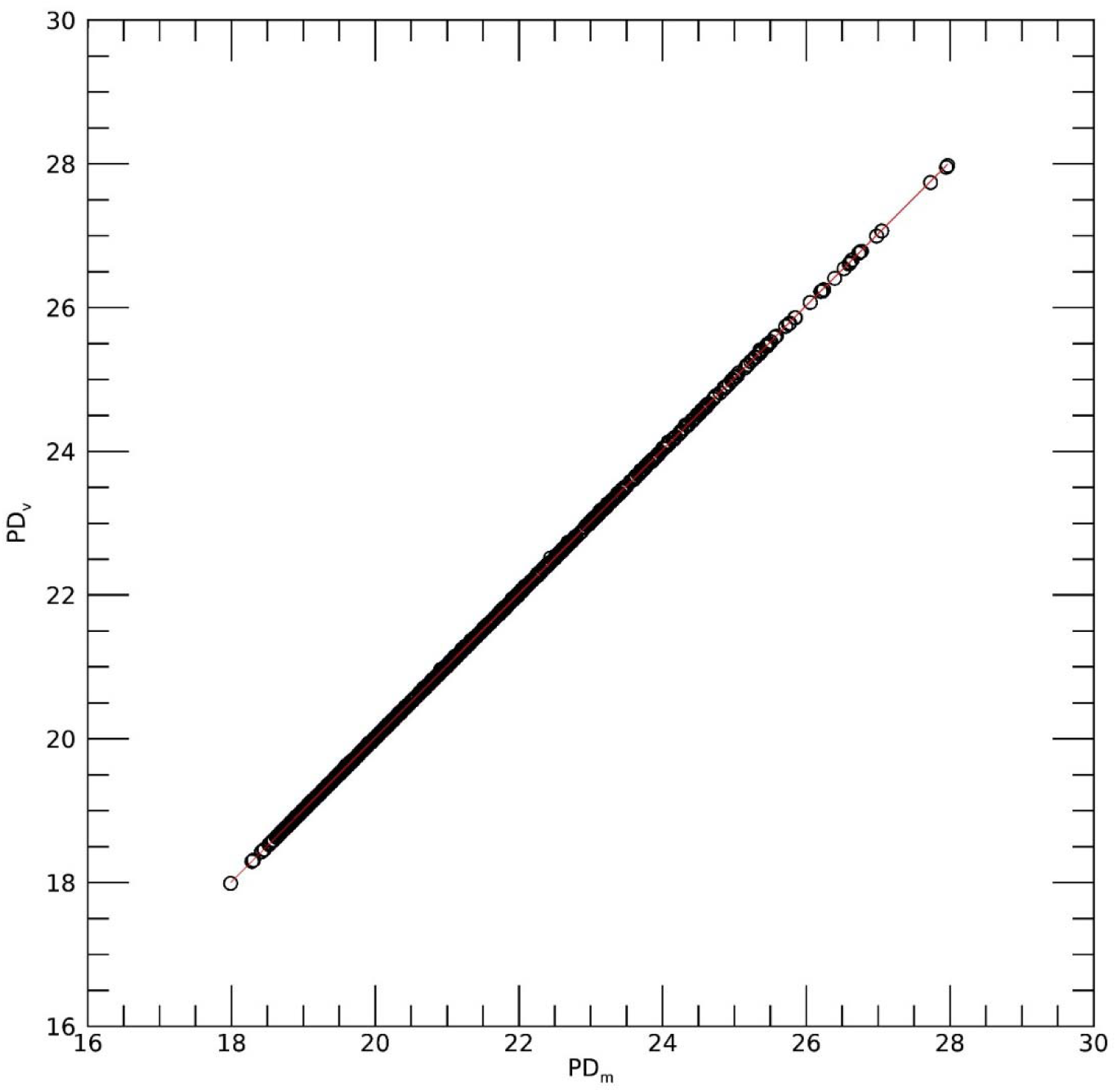
Regression analysis with PD_v_ as a function of PD_m_ This shows the scatter plot between the two measures (points) and regression line (solid red), The analysis gave: slope ≈ 1.002, intercept ≈ -0.0201, and linear correlation ≈ 1.0.

**Figure 9.**
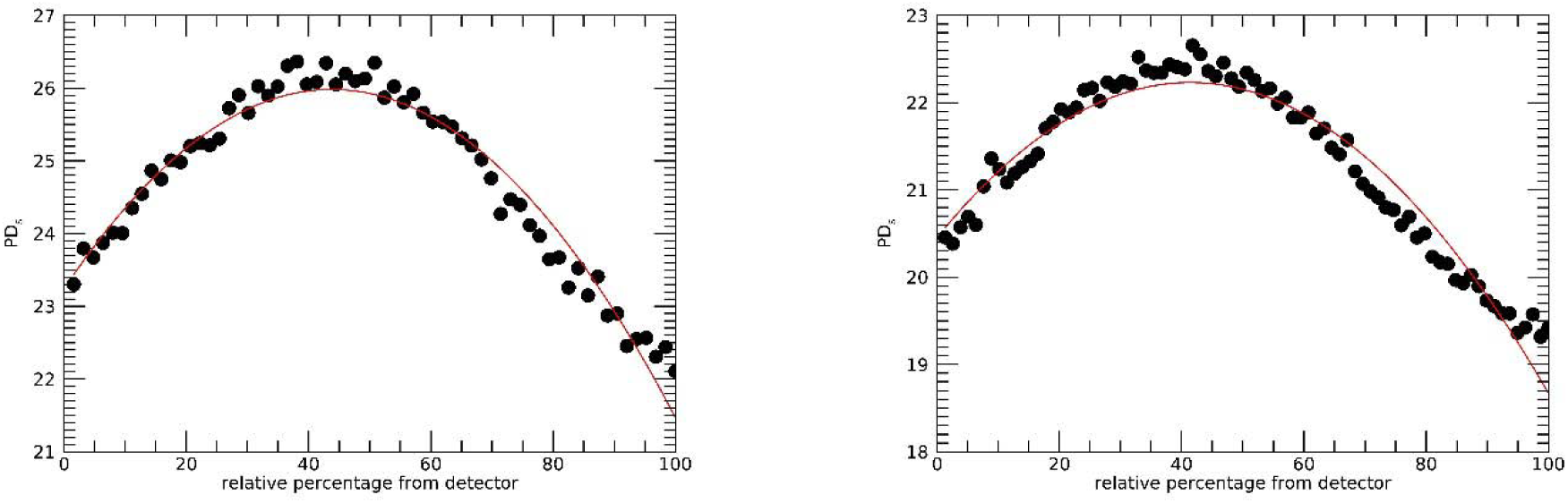
PD_s_ slice Profiles: this shows PD_s_ (y-axis) by slice number using the normalized distance (p) from the breast support surface (p on the x-axis) for illustration-1 (left) and illustration-2 (right). PD_s_ values per slice number (points) were fitted with a second-degree polynomial (solid). The slice distance increases as the distance increases from the breast support surface (left side of each plot).

**Figure 10.**
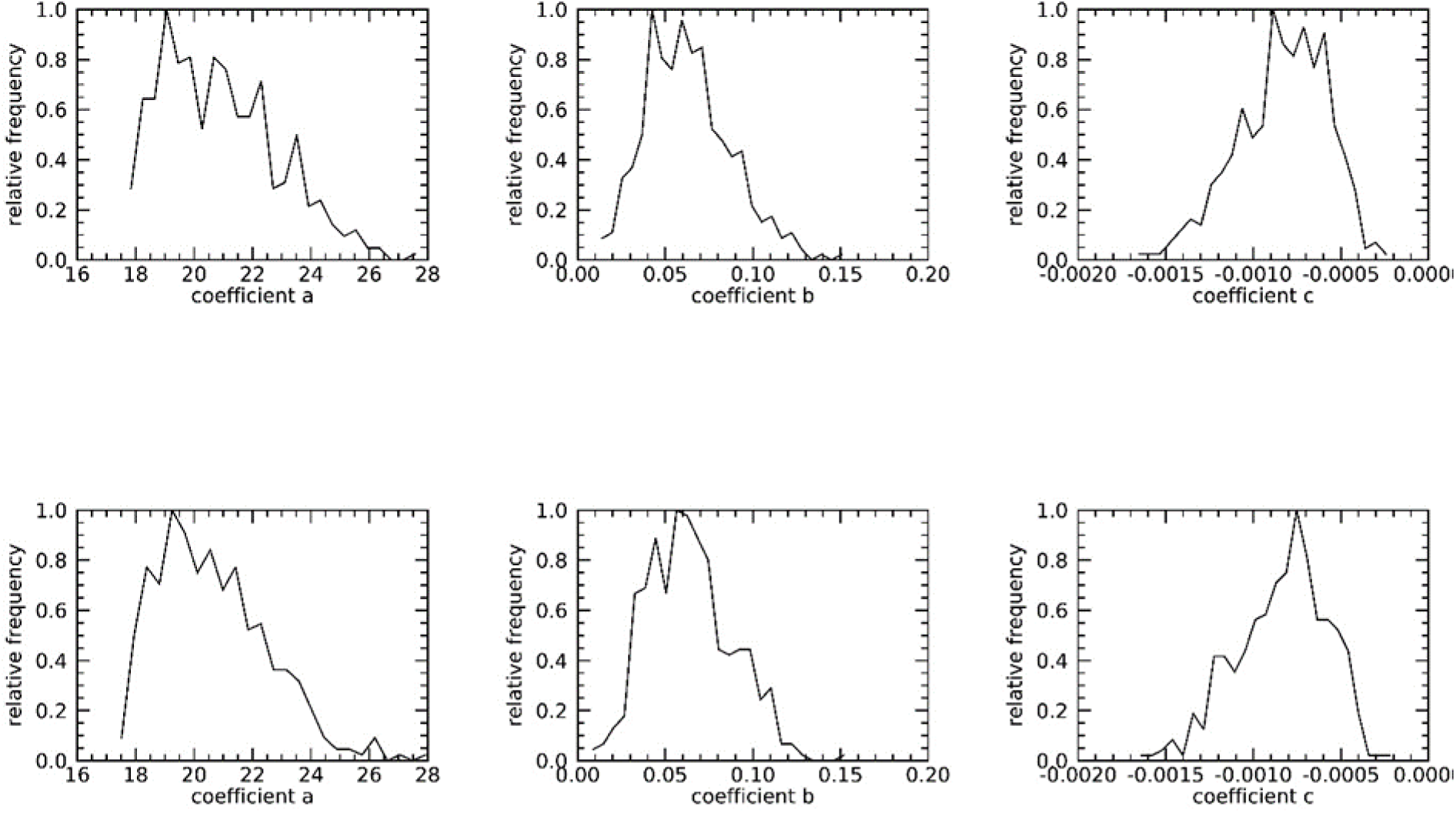
DBT Volume Slice 2^nd^ Degree Polynomial Coefficient Histograms: these show the normalized histograms for the fit-coefficients, (a,b,c), separated by cases (top-row) and controls (bottom-row).

### 3.4. Breast Cancer Risk Associations

Breast cancer associations are shown in Table 3. PD_syn_ was significant without noise multiplication [OR = 1.30 (1.08, 1.55)] and increased marginally with multiplication [OR =1.44 (1.18, 1.75)]. PD_v_ also provided a significant association with breast cancer [OR = 1.43 (1.18, 1.72)] that matched those of PD_m._ Isolating the analysis to single slices within the DBT volume produced significant associations: OR = 1.42 (1.17, 1.72) for the central slice and OR = 1.47 (1.21, 1.78) from the slice with the largest PD_s_ finding. The mean of the pixel values from the DBT volume (m_vol_) was also significant [OR = 1.31 (1.09, 1.57)] and very similar to those of PD_syn_ (without noise multiplication), and to that of mean from the C-View pixel values (m_syn_): OR = 1.29 (1.10, 1.52). Neither the total dense volume (D_v_) or the standard deviation of the pixel values within the volume (v_vol_) produced significant associations. We also investigated the differences between the maximum PD_s_ slice finding with the minimum finding from both the breast support surface (slice-1) and from the compression paddle (slice-100) as two different measures (not shown); these provided significant breast cancer associations that were similar to that of m_syn_ or m_vol_. When comparing the unadjusted models with adjusted models in Table 3, the ORs for the BD measurements shifted considerably. Therefore, we investigated the correlation between BMI and the three main PD_a_ applications by first removing BMI outliers giving: R = [PD_syn_, PD_v_, PD_m_] = [-0.43, -0.33, -0.33]. Although this range of correlation is weak, it explains the confounding influence of BMI on BD measurements; the risk of breast cancer increases as either BD or BMI increases while these two factors move in opposition.

**Table 3.**
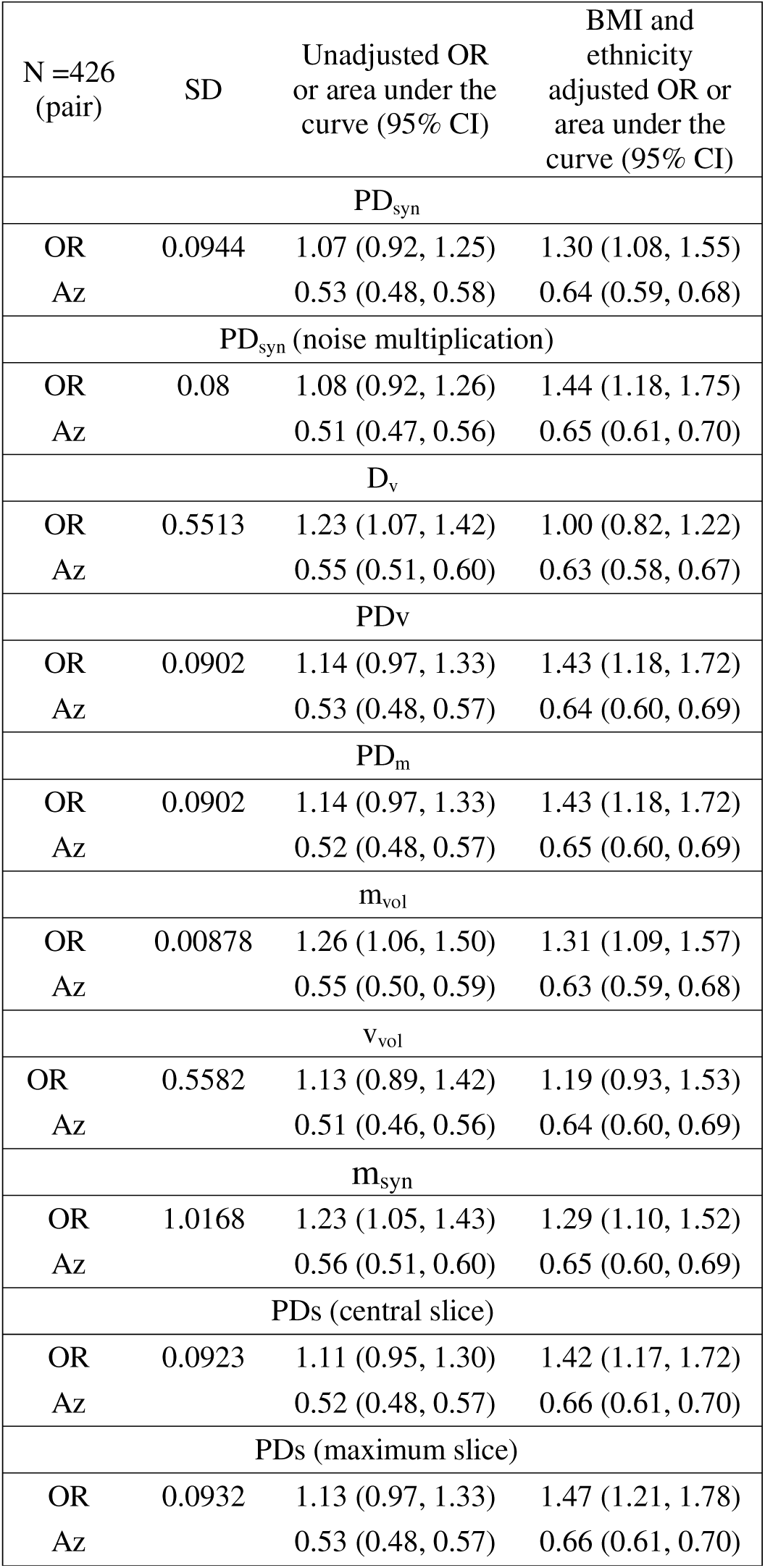
Conditional Logistic Regression Modeling Results: this table gives the odd ratios (ORs) with 95% confidence intervals (CIs) parenthetically for each model; image measures were log-transformed. The standard deviation (SD) is provided for each log-transformed distribution. The area under the receiver operating characteristic curve (Az) is given for each model with CIs parenthetically. From top to bottom: PD_syn_ is from C-View analysis; D_v_ is the dense volume; PD_v_ is volumetric PD; PD_m_ is the mean of the PD_s_ findings (mean of the volume slices); m_vol_ is the mean of the pixel values within the DBT volume (no processing); v_vol_ is the standard deviation of the pixel values within the DBT volume (no processing); and m_syn_ is the mean of the C-view (no processing); PD_s_ (central slice) was determined from the central slice from the DBT volume at p ≈ 50; PDs (maximum slice) was determined from the DBT slice with the largest PD_s_ quantity at p ≈ 41.

The OR findings above were similar for PD_v_ and PD_syn_. For risk factor purposes, this implies analyzing the C-View images (i.e., the breast volume structure projected onto a plane with heavy processing) is not subordinate to analyzing the volume images. To investigate these measures further, we investigated their relationship linear regression. Figure 11 shows the scatter plot (points) and regression analysis (solid line) with these findings: slope ≈ 0.92, intercept ≈ -1.4, and R ≈ 0.93. Because the slope is close to unity, the intercept is not far from PD_v_ = 0, and the strong positive correlation, these measures are approximately on the same scale and similar. Although the variation between these two measures increases as the respective measures increase, these findings explain the OR similarities found above.

**Figure 11.**
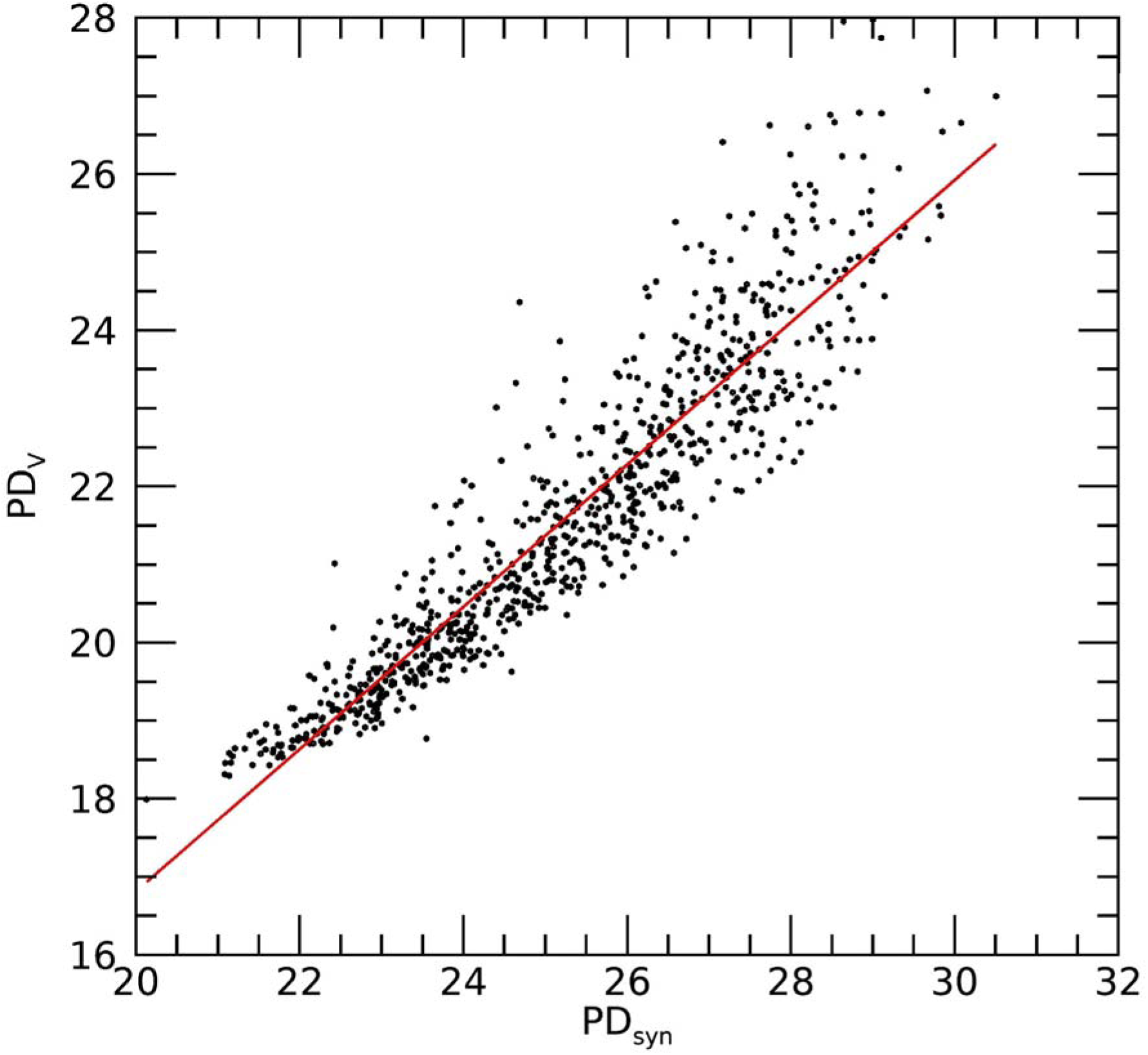
Regression analysis with PD_v_ as a function of PD_syn_: This shows the scatter plot between the two measures (points) and regression line (red-solid), The analysis gave: slope 0.92 with standard error ≈ 0.01, intercept ≈ -1.4 and R ≈ 0.93.

## 4. Discussion and Conclusions

We have provided an in-depth analysis of DBT data relative to an automated PD type breast density measure (here we refer to all measures in this study as PD for simplicity). First, the study provided evidence that the thresholds for the detection algorithm could be predetermined in advance by using noise multiplication. Qualitatively, labeled images with noise multiplication appeared superior, although ORs were only marginally larger. The volumetric PD measure was equivalent to the average taken across the DBT slice images, which agrees with the derivations that show the two measures are approximately equivalent. Three other findings were notable as well: (1) the DBT slice with the largest PD finding was offset considerably from the central slice, (2) PD from the central slice, from the slice with maximum PD finding, or from the C-View provided ORs similar to those from volumetric PD, and (3) the mean of the pixel values from the DBT volume slices or from the C-View images produced significant ORs without processing. The second finding also agrees with the mean slice results. The plots of slice PD profiles showed clusters of points (around 5-10 slices) with values close to PD_max_ about the curvature indicating why these isolated slices provided similar associations. We believe this is the first study to represent PD in this slice format. Breast cancer associations between the volume and the synthetic images were almost identical. This second finding indicates there is no benefit in terms of ORs derived from analyzing the volume; this applies to this method specifically but agrees with volumetric measures derived from 2D FFDM. Comparisons with other techniques are often not exact due to study design differences, in particular the sample size differences and model variations. Likewise, there is not an accepted convention for the standard deviation increment in the image measurement, which is distribution dependent for each measure. However, the ORs for PD found in this study agree with those determined previously [14, 44], are similar to those determined with volumetric measures [15] (from conventional 2D mammograms), and marginally less than a PD type measure applied to DBT [32]. Moreover, the third finding (i.e., significant ORs from pixel value means without PD processing) follows intuition in that larger pixel values represent elevated levels of dense breast tissue. We have found the variation in conventional 2D mammograms provided significant associations [6, 7, 15], which did not hold in this study for the DBT volume. D_v_ was not a significant risk factor but is the critical factor in the other measures; a significant measure is produced when normalizing this measurement by the total breast volume, which is in support of the probability conjecture. Thus, the study does provide evidence as to the nature of the traditional PD measurement and produced a related prescription for constructing a standardized synthesized 2D image.

There are some comments worth noting about this study. We analyzed a hospital-based population, where matching was used to account for case-control differences. Both the OR findings and summaries from Table 1 indicate this did not materially influence the outcomes. The PD derivations are general and apply to any like metric, whereas the findings in this report apply specifically to our automated approach. Although the results indicated that 2D and 3D measures from PD were similar, the study design establishes a template that could be used for investigating other measures such as texture. DBT is also shifting in technology. For instance, the manufacturer of the units used for this study is offering enhanced images from DBT, smaller pixel spacing, and interleaved slice spacing (increased). The noise field multiplication modification analyzed here offers potential to apply to images derived from evolving DBT advances. The results from this study will require verification in other populations and DBT technologies as well.

## Acknowledgements

This work was supported by the National Institutes of Health grants R01CA166269 and U01CA200464.

